# Optimizing Polygenic Scores for Complex Morphological Traits: A Case Study in Nasal Shape Prediction

**DOI:** 10.1101/2025.09.14.676081

**Authors:** Meng Yuan, Seppe Goovaerts, Nina Claessens, Jay Devine, Sara Becelaere, Isabelle Cleynen, Peter Claes

**Author notes:** Corresponding authors Meng Yuan; Peter Claes.

## Abstract

Polygenic scores (PGS) facilitate the prediction of an individual’s phenotype from their genotype. Typically, PGS methods apply regularization or use a clumping and thresholding (C+T) approach to handle SNP inclusion. To achieve good prediction accuracy, these approaches rely on effect size estimates from well-powered genome-wide association studies (GWAS). However, this is currently not feasible for morphological shape when phenotyped as univariate traits. Here, we introduce a novel framework to enhance polygenic prediction through three key components: (1)leveraging multivariate GWAS summary statistics for improved SNP selection, (2) defining genetically informative phenotypes, and (3) benchmarking PGS methods to select the optimal model. Our approach integrates multivariate GWAS, which performs an omnibus test against all phenotypic variables jointly with increased power. Specifically, our approach leverages *P* values from multivariate GWAS to improve SNP selection while maintaining the effect size estimates for the univariate trait under investigation, allowing the use of current PGS tools. We evaluated our proposed method for predicting 3D nasal morphology using a dataset of 52,896 individuals of European ancestry from the UK Biobank. Using the C+T method, SNP selection based on multivariate GWAS resulted in significantly improved phenotypic prediction (*P* = 9.74e-5) for eigen-shapes, with a mean variance explained of 3.88% (SD = 1.59%) compared to 2.02% (SD = 1.10%) using a traditional univariate approach in the test set (n = 2,896). We also tested whether heritability-optimized phenotypes were more predictable than eigen-shapes derived from principal component analysis (PCA). On average, with SNP selection based on multivariate GWAS using the C+T method, heritability-optimized phenotypes yielded greater predictive performance, with PGS scores explaining 2.72%-10.37% of phenotypic variance, compared to 1.05%-6.84% for eigen-shapes. Furthermore, benchmarking several PGS methods revealed that LDpred2 consistently achieved the best performance for predicting nasal morphology. Our results demonstrate that combining multivariate GWAS *P* values with optimized phenotypes and advanced PGS models leads to more accurate polygenic prediction for complex morphological traits.

## Introduction

Predicting an individual’s genetic predisposition for a complex disease or trait is a critical task known as polygenic scoring that has far-reaching scientific, ethical, and societal implications. For many genetic disorders, polygenic scores (PGS) show promise in early risk detection, stratification of patient groups, and predicting therapeutic responses [1], [2], [3], [4], [5]. PGS are typically calculated as the weighted sum of allele dosages of single nucleotide polymorphisms (SNPs), where trait-associated SNPs are usually identified in genome-wide association studies (GWAS). The weights reflect the magnitude of association between each SNP and the trait. Unfortunately, PGS cannot serve as stand-alone prediction tools, as they capture only a fraction of the genetic contribution, with other genetic and non-genetic factors also playing a significant role [6].

Complex morphological traits are particularly difficult to predict because their phenotypic variation is multipartite, and no common consensus exists to define phenotypes. Moreover, morphological traits are highly polygenic, shaped by many genetic effects that drive spatially and temporally dependent developmental processes acted on environmental context [7], [8], [9]. To date, the most comprehensive review on facial shape in European individuals estimated that 501 independent SNPs across 303 loci explain only 13.7% of the sample variance [10]. Another study focused on variants with <1% frequency found seven genes to be enriched for rare variants affecting facial shape [11]. In general, our current understanding of common and rare genetic variants is unlikely to yield an accurate reconstruction of the entire multivariate face due to this complexity. Optimizing a latent trait that is highly heritable based on a multivariate description and then predicting these distinct univariate facial traits, on the other hand, is a more feasible task that has not been fully explored.

Various methods have been proposed to calculate PGS, differing primarily in two key aspects: the selection of SNPs and the weights assigned to them [12]. The standard approach, clumping and thresholding (C+T), selects a set of approximately independent SNPs by applying linkage disequilibrium (LD) clumping to SNPs that exceed a predefined *P* value threshold [13], [14], [15]. Although computationally and conceptually simple, the C+T method uses only a subset of SNPs, ignoring many other SNPs along with their LD information, which limits prediction accuracy [16]. To address this, more sophisticated PGS methods have been developed to incorporate genome-wide SNPs and their LD structure. These methods typically perform shrinkage on all SNP effect sizes, leveraging commonly used regularization techniques (e.g., Lassosum [17], Lassosum2 [18]) or Bayesian shrinkage approaches (e.g., LDpred [19], LDpred2 [20], and PRS-CS [21]).

Recent comparative studies of PGS methods have shown that no approach is universally optimal, as prediction accuracy depends on trait-specific genetic architecture, the statistical power of GWAS, and ancestry composition [22], [23]. While sufficiently powered GWAS are essential for accurate PGS, this remains currently unachievable for facial traits when phenotyped univariately, necessitating alternative strategies. Previous multivariate GWAS of facial shape have uncovered more genomic loci than their univariate GWAS counterparts [10]. Open-ended multivariate GWAS improve genomic discovery because they leverage cross-trait genetic covariance, while univariate GWAS ignores this. In addition, they reduce the multiple-testing burden compared to analyzing all genetically correlated traits separately [24].

Here, we investigate the optimization of three distinct components to enhance PGS prediction of complex morphological traits: SNP selection, phenotype definition, and the PGS model. Three-dimensional (3D) nose morphology, an easy to recognize and heritable [25] part of the human face, is used as a case study. First, we propose the use of multivariate GWAS *P* values to improve SNP selection, leveraging their ability to capture a variant’s association with multiple traits simultaneously. We retain univariate effect size estimates for PGS construction, ensuring compatibility with existing tools designed for single-trait prediction. Second, rather than predicting the full nasal shape, we focus on distinct nasal features, evaluating how different phenotype definitions influence predictability. To this end, we compare three types of nasal phenotypes: (1) widely used traditional anthropometric measures (i.e., inter-landmark distances);eigen-shapes derived from principal component analysis (PCA); (3) heritability-enriched traits generated through optimization algorithms [26]. Third, we benchmark several PGS methods, including PRSice-2 [15], LDpred2 [20], Lassosum2 [18] and PRS-CS [21], and identify the best-performing model for nasal trait prediction.

## Results

In this work, we considered three key components in PGS construction, as illustrated in Fig. 1A: (1) phenotype definition, (2) selection of SNPs, and (3) weight assignment strategies in PGS methods. Nose morphology data (n = 52,896) was sourced from the UK Biobank [27] and extracted from full head MRI images (see Methods section). From this dataset, 45,000 samples were used as training data to perform discovery GWAS and primary PGS model building. A separate set of 5,000 samples was used as validation data for hyperparameter fine-tuning (Fig. 1B). Finally, 2,896 samples were used as independent test data to evaluate prediction performance (Fig. 1C). Prediction performance was assessed by fitting a linear model where PGS scores were used to predict the original phenotypes, with R^2^ (coefficient of determination) quantifying the proportion of phenotypic variance explained. Higher R^2^ indicates better prediction.

**Figure 1.**
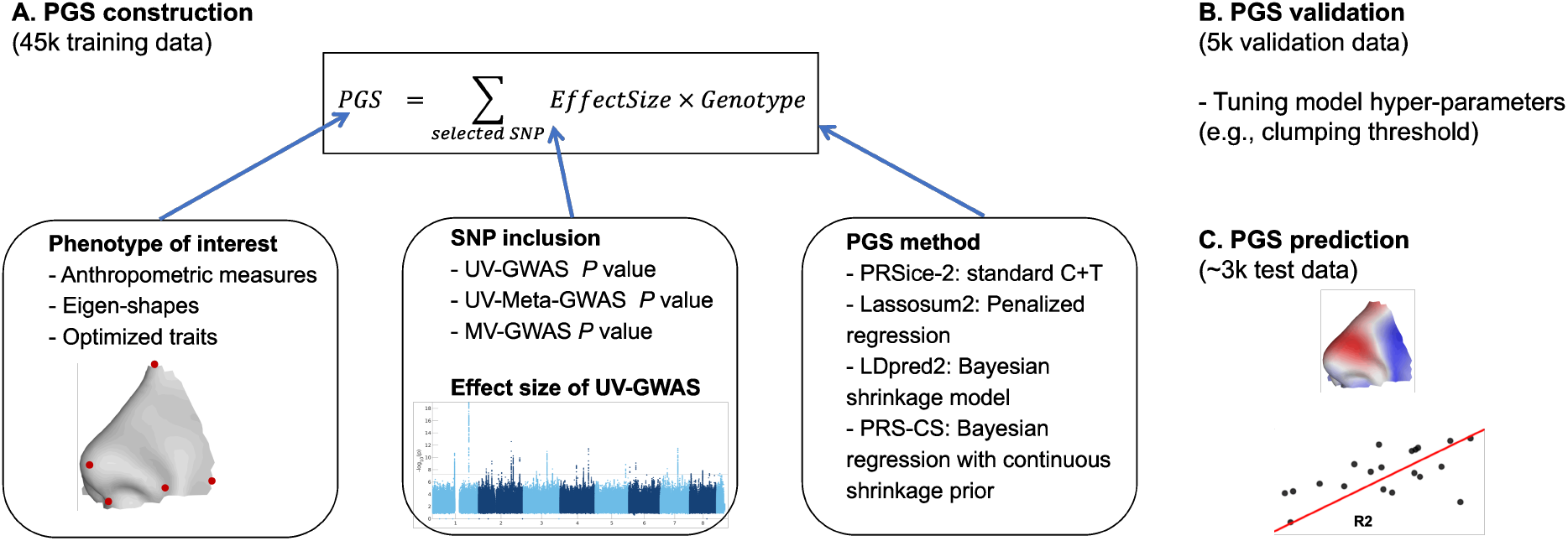
PGS workflow.

1. Phenotype definition: The phenotype of interest was defined in three ways: (1) anthropometric measures (10 linear distances among 5 sparse landmarks as described previously in [25], see Supplementary File 2); (2) 21 eigen-shapes derived from applying PCA to dense landmarks, or (3) 21 heritability-enriched traits obtained by training a genetic algorithm (GA) model to identify directions in feature space with high heritability [26] (see Methods).
2. SNP inclusion: SNPs associated with the phenotype of interest are typically identified through univariate GWAS (UV-GWAS; a separate GWAS for each trait), where both SNP selection and weight estimation rely on the same discovery GWAS. In this study, we used univariate effect size estimates of UV-GWAS as weights to ensure compatibility with existing PGS tools and explored alternative SNP inclusion strategies. First, we used meta-analyzed univariate GWAS (UV-Meta-GWAS) *P* values for SNP selection, which aggregates univariate GWAS results within each phenotype category (see Methods). Second, we evaluated *P* values from multivariate GWAS (MV-GWAS) for SNP selection. The MV-GWAS was conducted on eigen-shapes using canonical correlation analysis (CCA), a well-established method that has successfully identified genomic loci associated with craniofacial morphology by performing an omnibus test against all phenotypic variables jointly with increased power [28].
3. Methods benchmarking: Some studies [12], [22], [23] have compared various PGS methods for other types of traits; however, none have benchmarked these methods for complex morphological traits. Therefore, we utilized a PGS toolbox (https://github.com/SaraBecelaere/STREAM-PRS) which includes the C+T approach implemented in PRSice-2 [15], and several effect size shrinkage methods such as LDpred2 [20], Lassosum [17], Lassosum2 [18] and PRS-CS [21]. Detailed descriptions of training and hyperparameter settings for each method are provided in Methods section.

Fig. 2 compares prediction performance under different SNP inclusion strategies using the C+T approach (numerical details in Supplementary File 1, Table S1). A consistent pattern emerged across all phenotype groups: MV-GWAS yielded the best results, followed by UV-Meta-GWAS, while standard UV-GWAS performed the least effectively. A substantial improvement was observed for heritability-enriched traits and eigen-shapes using MV-GWAS for SNP selection (mean variance explained of 5.86% [SD=1.86%] vs. 3.51% [SD=1.55%] and 3.88% [SD=1.59%] vs. 2.02% [SD=1.10%], respectively). Meanwhile, inter-landmark distances exhibited a relatively moderate gain (4.71% [SD=0.98%] vs. 3.15% [SD=1.22%]). When comparing phenotype definitions using the simple C+T approach and MV-GWAS, heritability-enriched traits were more predictable than eigen-shapes (median R^2^: 5.57%; range: 2.72-10.37% vs. median 4.39%; range: 1.05-6.84%; *P* = 1.3e-3). Inter-landmark distances demonstrated exhibited a moderate level of predictability (median: 4.66%; range: 3.40-6.02%).

**Figure 2.**
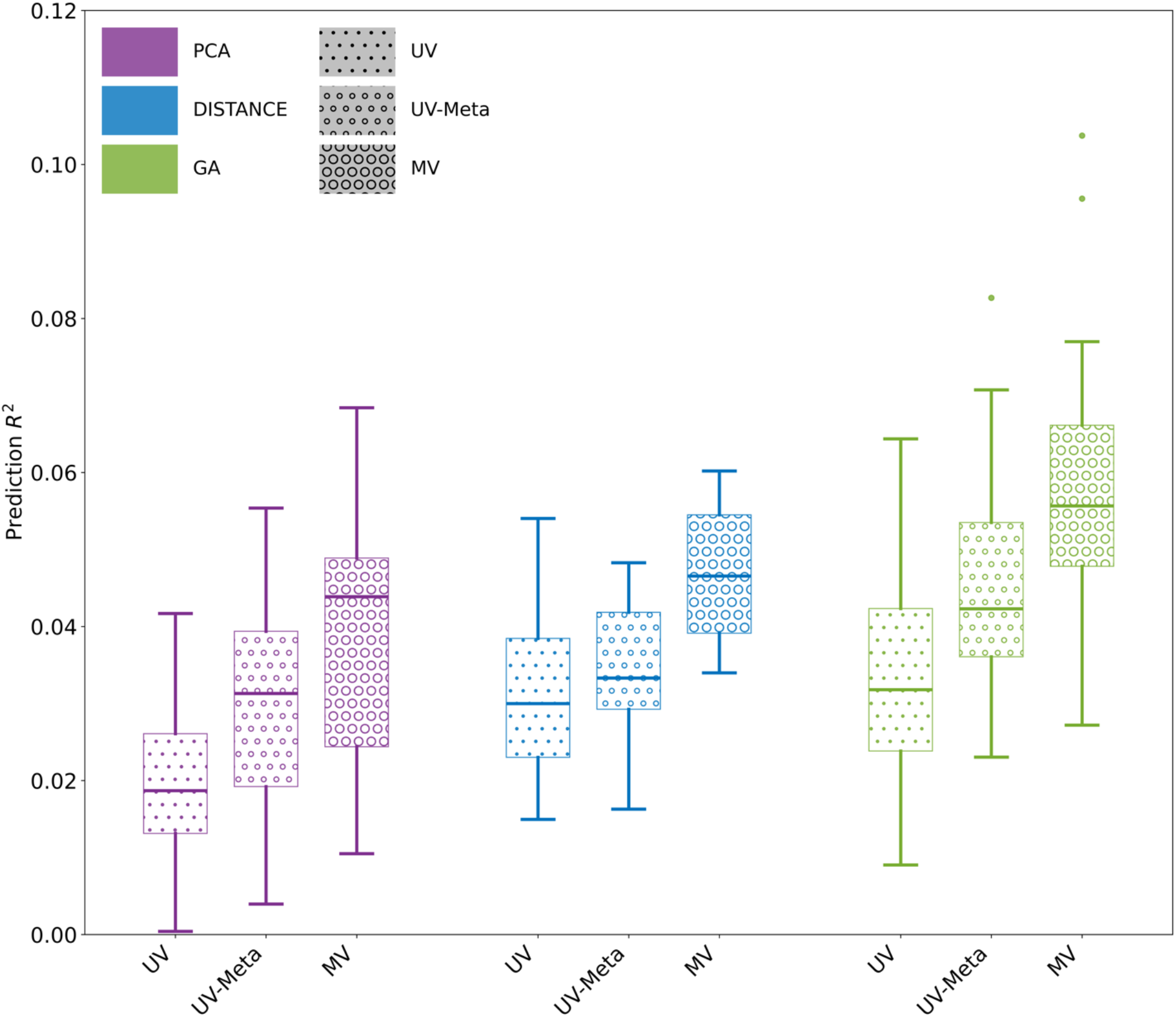
Comparison of prediction performance using the C+T approach.

After confirming that multivariate GWAS *P* values result in more optimal SNP selection, we compared PGS approaches under this scheme (Fig. 3). LDpred2 consistently showed relatively higher performance across all phenotypes. For inter-landmark distances, LDpred2 achieved a higher mean explained variance (7.57%) than PRSice-2 (4.71%; *P* = 8.5e-3) and PRS-CS (5.02%; *P*

**Figure 3.**
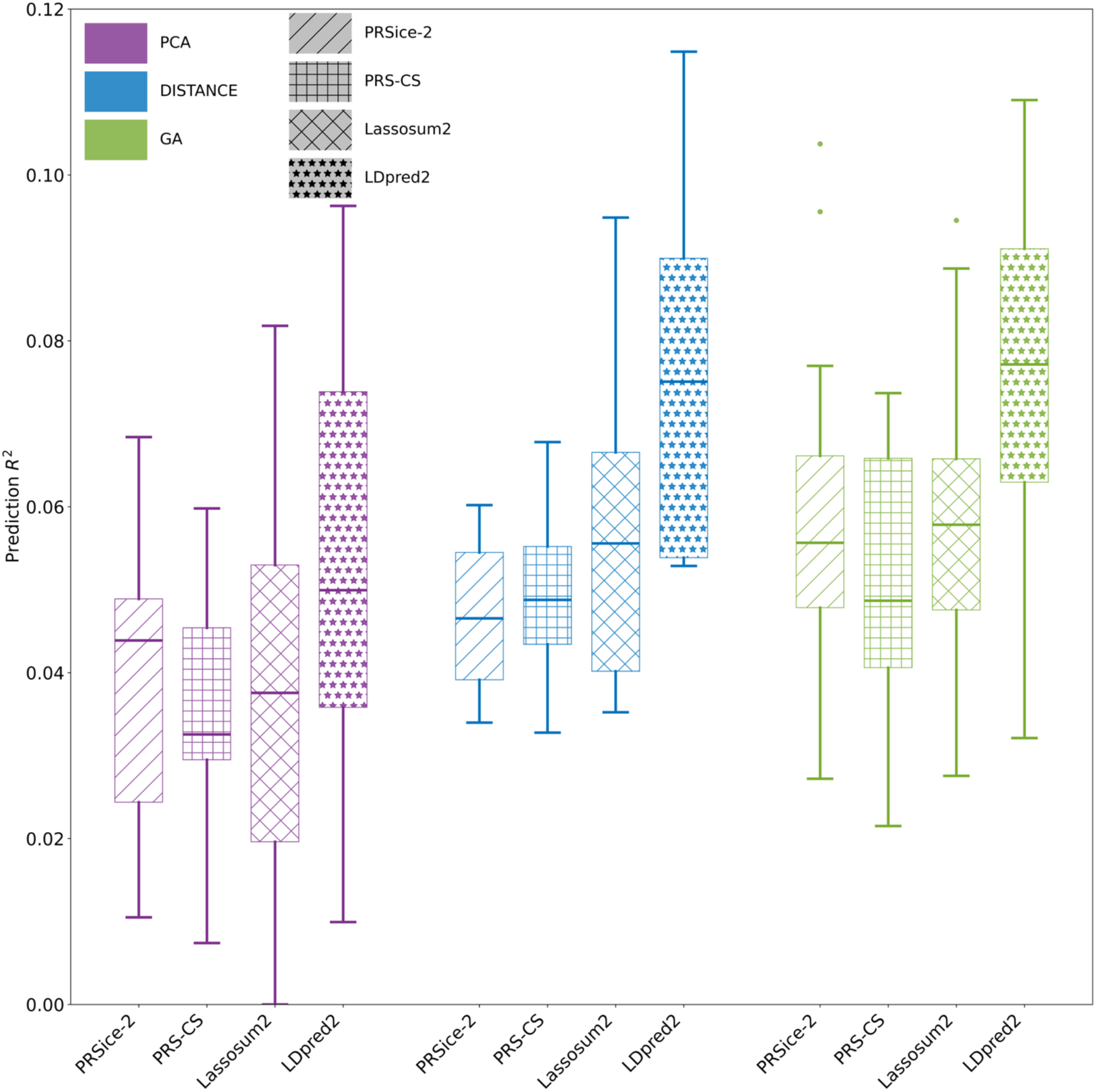
Comparison of prediction performance using different phenotyping methods and PGS approaches with multivariate GWAS *P* values for SNP inclusion.

= 1.85e-2) and was not significantly different from Lassosum2 (5.63%; *P* = 1.59e-1); all *P* values are Bonferroni-corrected for three tests. When comparing phenotype definitions using LDpred2, heritability-enriched traits and inter-landmark distances showed similar performance, both surpassing PCA-based eigen-shapes predictions (details in Supplementary File 1, Table S2).

Fig. 4 illustrates the prediction performance for 10 representative traits, including linear distances, the first 10 eigen-shapes, and the first 10 heritability-enriched traits (visualizations of all nasal traits are in Supplementary File 2). The highest phenotypic variance explained was observed for the linear distance between the nasion and cheek points (R^2^ = 11.5%, Fig. 4A, column 7). Similarly, a heritability-enriched trait (Fig. 4C, column 5), characterized by a larger, rounded, and pointing nose tip, also exhibited high predictability (R^2^ = 10.3%). Notably, another heritability-optimized trait (Fig. 4C, column 10), which effectively captures fine-scale morphological features (particularly alar crease curvature), achieved high predictive performance (R^2^ = 10.9%). In contrast, a similar linear distance measure (Fig. 4A, column 10) showed relatively low performance (R^2^ = 5.3%), demonstrating that linear measures alone cannot adequately represent complex curvature features.

**Figure 4.**
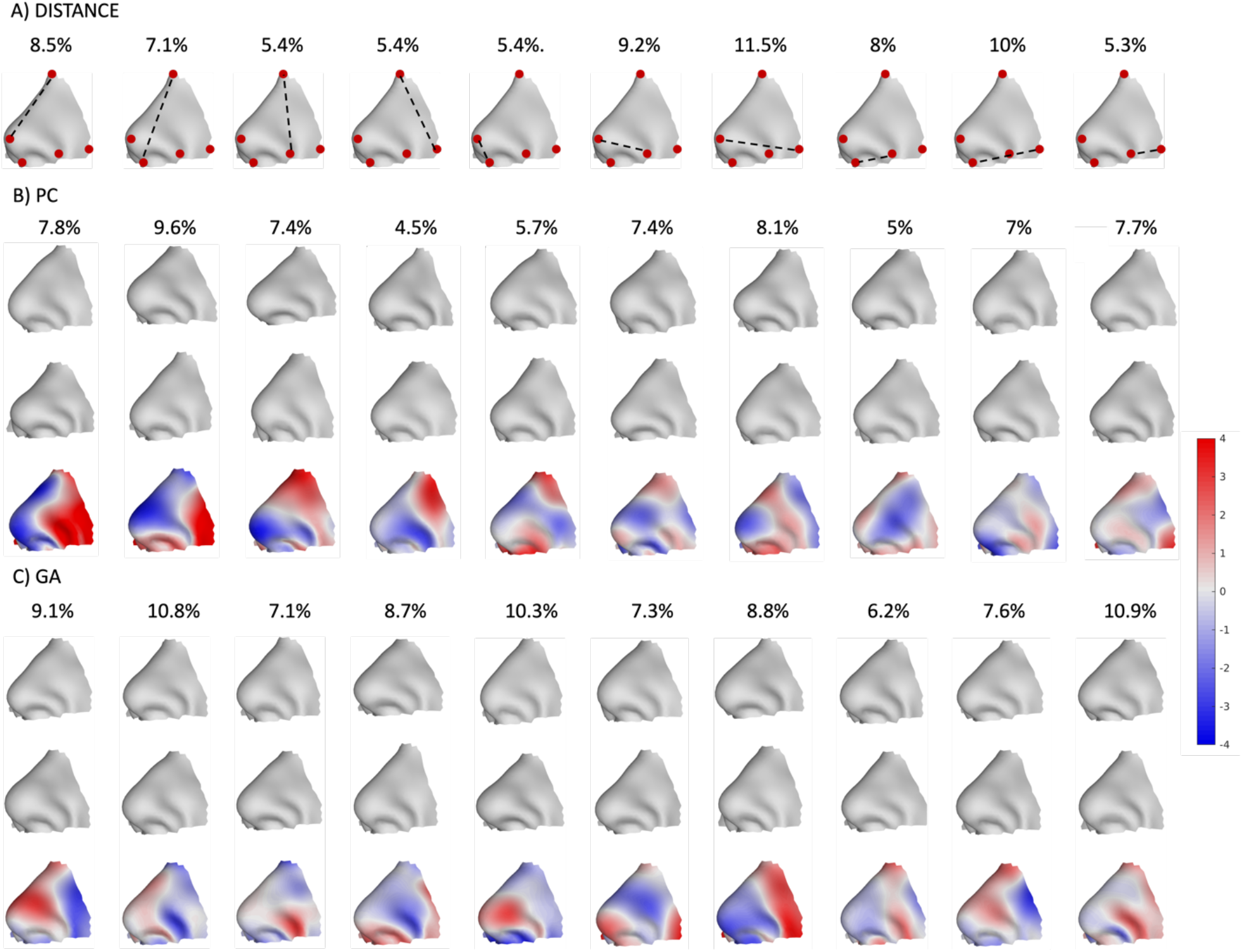
Visualization of nasal traits. (A) The first panel displays the 10 linear distances alongside their corresponding prediction performance. (B) The second panel displays the first 10 principal components (PCs). Shape variation is illustrated in gray, showing the range from the mean to plus and minus 3 times the standard deviation (SD) along each PC, positioned vertically with respect to each other (top + 3SD, and bottom -3D). Colored bars represent differences (in mm) between these two opposite deviations from the mean. (C) Similarly, the third panel presents the first 10 heritability-enriched traits.

## Discussion

In summary, we proposed a framework to optimize PGS construction for complex morphological traits through three key components: (1) leveraging multivariate GWAS summary statistics to enhance SNP selection, (2) defining genetically informative phenotypes by exploiting the inherent multivariate nature of morphological traits, and (3) benchmarking PGS methods to identify the best-performing model. We evaluated the framework using a large dataset from the UK Biobank and found that incorporating MV-GWAS *P* values significantly improved PGS prediction accuracy compared to standard UV-GWAS. Among phenotype definitions, heritability-enriched traits and inter-landmark distances yielded better performance than eigen-shapes. Furthermore, comparisons across PGS models revealed that LDpred2 consistently outperformed other methods in predicting nasal morphology.

A key contribution of our study is the use of open-ended MV-GWAS summary statistics to refine SNP selection in PGS development. MV-GWAS identified more significant associations than UV-GWAS, capturing all genomic loci detected by UV-GWAS while also uncovering additional ones [10]. This is particularly advantageous for complex traits like the human face, where genetic variants often affect multiple facial traits simultaneously, each with small effect sizes. By accounting for trait dependencies, MV-GWAS enhances statistical power to identify pleiotropic SNPs. As expected, UV-Meta-GWAS, which is another way to boost power, performed intermediately, falling between UV-GWAS and MV-GWAS. From a computational standpoint, integrating MV-GWAS *P* values into existing PGS toolboxes is straightforward, requiring only separate inputs for SNP selection (i.e., *P* values) and weight estimation (i.e., allele effect sizes). Given its superior performance in all comparisons, we strongly recommend using MV-GWAS for selecting candidate SNPs in PGS calculation for complex traits, especially for morphological traits, such as facial structure, cranial vault shape, or brain morphology.

The multivariate nature of morphological traits provides an opportunity to optimize latent traits for heritability. Accordingly, heritability-enriched traits demonstrated higher prediction performance than PCA-based eigen-shapes across all PGS methods. This aligns with previous findings [26], [29], as these phenotypes were explicitly optimized to capture genetically informative features of nasal shape, making them better suited for genotype-phenotype analyses. While PCA remains a valuable tool for extracting features from high-dimensional data, its derived features are driven by statistical variance explained which may not correspond well to heritability. Supporting this, previous studies [26], [29] have demonstrated that heritability-enriched phenotypes exhibit higher SNP-heritability and more effectively identify relevant genomic loci compared to eigen-shapes.

When using the C+T approach, heritability-enriched traits yielded relatively higher predictive accuracy than inter-landmark distances. However, both approaches achieved comparable performance with optimal PGS models. Inter-landmark distances are defined based on anatomical and biological prior knowledge [30], [31], and a previous study [25] has shown their high SNP-heritability, indicating strong alignment with genetically determined shape variation. A limitation of this approach, however, is that the genetic findings are constrained by the available landmarks. This is particularly evident in nasal shape analysis (Fig. 4), where subtle changes in complex morphology (e.g., curvature) cannot be adequately represented by only five landmarks and their inter-landmark distances. In contrast, heritability-enriched traits are optimized from dense landmarks and can capture highly heritable, localized regions with greater precision. Ultimately, this represents a trade-off: inter-landmark measures are simpler to define but require advanced PGS models (e.g., LDpred2) for accurate prediction, whereas heritability-enriched traits require training sophisticated optimization algorithms to extract features from dense landmarks but achieve strong prediction performance even with basic PGS methods (e.g., C+T approach).

A comparison of four PGS methods (PRSice-2 [15], LDpred2 [20], Lassosum2 [18] and PRS-CS [21]) revealed that LDpred2 achieved the highest prediction accuracy overall. While some studies [12],

[20] suggest that PGS methods which more formally model trait-specific genetic architecture (e.g., LDpred2, PRS-CS) outperform simpler approaches like C+T approach, we found that the C+T method remained competitive for nasal morphology. This may be due to the highly polygenic nature of nasal traits, and thresholding methods generally outperform PRS-CS for such traits [22]. Moreover, optimal parameter settings for PGS methods that account for the genetic architecture of specific traits could further enhance the prediction performance. Computationally, the C+T approach (as implemented in PRSice-2) was the most efficient, taking ∼20 minutes per univariate trait on a single CPU to first generate SNP weights based on the discovery sample, then apply them to the validation and testing samples. In contrast, PRS-CS required ∼6 hours, while LDpred2 and Lassosum2 (jointly implemented in [32]) took ∼3 hours in total. To improve prediction accuracy, we recommend applying multiple PGS methods, selecting the best-performing model, and using a grid search to tune hyperparameters.

While our framework advances PGS construction for nasal traits, and by extension facial traits, the practical utility of current models remains limited. Even the best-performing models explained only ∼10% of phenotypic variance, a level unlikely to be meaningful for most real-world applications in forensic, anthropological, and clinical research. This limitation arises because PGS captures only the additive genetic component of a trait, excluding non-genetic factors and gene– environment interactions; thus, its predictive accuracy is inherently bounded by heritability [6], [33]. Additionally, GWAS do not account for transcriptomic or proteomic variation further limiting their explanatory power [34]. Translating PGS into practice also faces significant clinical and social hurdles, as naive implementations risk introducing severe bias and misinterpretation [35], [36]. Given these challenges, it may be more practical to focus on predicting specific, distinctive traits rather than attempting to predict the entire morphological shape.

A limitation of our study is its focus on individuals of European ancestry. Although increased diversity introduces statistical challenges, methods like PRS-CSx [37] can effectively couple genetic effects across populations through a shared prior. Studies have demonstrated that training PGS models in ancestrally diverse cohorts improves the weight estimation of genetic variants, particularly for variants with higher frequencies in non-European populations [22], [23]. For example, a multi-ancestry GWAS meta-analysis of inter-landmark distance-based facial features explained 1.61%–7.50% of the phenotypic variance [38]. Their performance is comparable to ours, despite being based on a smaller sample size that included 9,674 East Asians and 10,115 Europeans. This suggests that incorporating diverse ancestries may enhance PGS model performance, or the relatively high phenotypic variance explained is partially driven by population-level (ancestry-related) structure in addition to individual-level differences. Future research could explore multi-ancestry PGS approaches to enhance prediction accuracy and understand the properties of our proposed strategies within and across ancestries.

In conclusion, we introduced a framework to optimize PGS construction for complex morphological traits by enhancing SNP selection through the integration of MV-GWAS summary statistics, genetically informative phenotype definitions, and PGS model selection. When applied to the prediction of nasal morphology, this framework identified distinctive traits with notably higher predictability. Our findings offer valuable insights for future PGS-based predictions, particularly for multivariate traits or, more broadly, sets of genetically correlated traits.

## Materials and methods

### Dataset

Our study utilized data from the UK Biobank [27], which comprises genetic and head MRI data from approximately 60,000 participants in the UK. We restricted our analysis to unrelated individuals of European ancestry. Ancestry assignment for each participant was based on the Pan-UK Biobank Project (https://pan.ukbb.broadinstitute.org/). Related individuals were identified using the KING-robust [39] kinship estimator at a threshold of 0.0442 (third-degree relatives) followed by random selection of one individual per related group. Genotype imputation procedures for the UK Biobank have been described in [27]. After imputation, we applied quality control filters to retain variants with an imputation INFO score > 0.3, minor allele frequency (MAF) > 0.01, genotyping missingness rate < 5%, and Hardy–Weinberg equilibrium (HWE) *P* value > 1 × 10-6. Additionally, individuals with more than 5% missing genotype data were excluded from the analysis. Phenotype processing, including 3D image quality control and sparse/dense landmarking, was performed as described in [Goovaerts et al, in preparation]. The final dataset included 8,922,008 SNPs and 52,896 individuals. We partitioned the data into 45,000 samples for discovery GWAS and primary PGS model training, 5,000 for validation and hyperparameter tuning, and 2,896 for evaluating prediction performance.

### Phenotyping methods

The phenotype of interest included anthropometric traits, eigen-shapes, and heritability-enriched traits, with detailed methodologies described in previous work [25], [26]. Briefly, we focused on 5 anatomical nasal landmarks and computed 10 inter-landmark Euclidean distances between landmarks (Supplementary File 2). Besides sparse landmarks, we also used dense landmark configurations, represented as a 3D matrix of dimensions N (number of shapes), L (7,160 quasi-landmarks), and 3 (x, y, z coordinates). After mean-centering and reshaping the landmarks into a 2D matrix, we applied low-rank singular value decomposition (SVD). To retain meaningful variation, we combined PCA with parallel analysis [40], [41], yielding 21 eigen-shapes that captured 98.21% of nasal shape variation. While eigen-shapes are widely used, they are unsupervised and may not align with genetically relevant phenotypic axes. Therefore, we applied a third approach optimizing phenotype extraction for genetic analyses. As described in [26], we applied PCA to construct a lower-dimensional feature space encoding complex shape variations. Then, we employed an optimization algorithm, more explicitly a genetic algorithm (GA), to identify directions or traits in this space with high SNP-heritability (training details and hyperparameter settings are in Supplementary File 3). SNP-heritability was computed via GREML [42], [43] based on unrelated individuals using SNPlib toolbox [44] (https://github.com/jiarui-li/SNPLIB).

### Genome-wide association analysis

For UV-GWAS, we performed linear regression (function ‘regstats’ from Matlab 2022b) under an additive genetic model (SNP dosages: 0, 1, 2), adjusting for covariates (sex, age, age-squared, height, weight, and the first ten genomic ancestry axes), prior to the regression. To obtain *P* values for UV-Meta-GWAS, we took the lowest *P* value for each SNP across all traits within the same phenotype group.

For MV-GWAS, we treated the 21 eigen-shapes as a unified representation of multivariate shape variation. Using CCA (function ‘canoncorr’ from Matlab 2022b), we identified the linear combination of these 21 components that maximally correlated with the SNP dosage in the discovery cohort. Prior to CCA, we adjusted for the same set of covariates used in the UV-GWAS.

### PGS method benchmarking

PGS method benchmarking was performed using STREAM-PRS toolbox as described in [Becelaere et al, in revision], which incorporates the C+T approach implemented in PRSice-2 [15], and shrinkage-based methods LDpred2 [20], Lassosum2 [18] and PRS-CS [21]. For the C+T method, we tested a series of *P* value thresholds: 5e-8, 1e-7, 1e-6, 1e-5, 1e-4, 0.001, 0.01, 0.02, 0.05, 0.1, 0.2, 0.3, 0.4, 0.5, and 0.6. Clumping was performed using a 250 kb window and an LD R^2^ threshold of 0.1. For LDpred2 (jointly implemented with Lassosum2 in [32]), we applied the LDpred2-inf, LDpred2-grid, and LDpred2-auto models. When running LDpred2-grid, we tested a grid of hyperparameters, with the proportion of causal variants p from a sequence of 10 values from 1e-6 to 1 on a log-scale and the heritability coefficient h^2^ = (0.3, 0.7, 1, 1.5). For Lassosum2, we used a grid of hyperparameters: nLambda = 20, lambdaMinRatio = 0.01, and delta = (0.001, 0.05, 1). For PRS-CS, we tested the global shrinkage parameter phi = (1, 0.1, 0.01, 0.001, 1e-4, 1e-6). For all PGS models, optimal hyperparameters were selected using a validation set. Prediction performance was evaluated by fitting a linear model in which the PGS scores were used to predict the original phenotype, with R^2^ (coefficient of determination) quantifying the proportion of phenotypic variance explained.

We considered three key components in PGS construction: phenotype definition, SNP inclusion, and weight assignment strategies in PGS models. (A) A total of 45,000 samples were used as training data to perform discovery GWAS and to build the PGS model; (B) 5,000 samples were used as validation data for hyperparameter fine-tuning; and (C) 2,896 samples were used as an independent test set to evaluate prediction performance.

We evaluated three categories of phenotypes: (1) anthropometric measures (i.e., Distance), (2) eigen-shapes derived from principal component analysis (PCA), and (3) heritability-enriched traits obtained by training a genetic algorithm (GA). Three SNP inclusion strategies were tested:

(1) standard univariate GWAS (UV-GWAS); (2) SNPs identified based on meta-analyzed univariate GWAS (UV-Meta-GWAS), which aggregates univariate GWAS results within each phenotype category; and (3) multivariate GWAS (MV-GWAS), where we treated the 21 eigen-shapes as a unified representation of multivariate shape variation and performed GWAS using canonical correlation analysis (CCA).

We evaluated three categories of phenotypes: (1) anthropometric measures (i.e., Distance), (2) eigen-shapes derived from principal component analysis (PCA), and (3) heritability-enriched traits obtained by training a genetic algorithm (GA). Four PGS methods were compared: the C+T approach (implemented in PRSice-2), PRS-CS, Lassosum2 and LDpred2.

## Ethical approval

The committee/institutional research board of UK Biobank gave ethical approval for collection of the UK Biobank data (https://www.ukbiobank.ac.uk/learn-more-about-uk-biobank/about-us/ethics). Approval to use UK Biobank at an individual level in this work was obtained under application no. 88320. Local ethics approval at the KU Leuven, Belgium, was provided under PRET G-2022-5272.

## Data availability

All the data and detailed information for the UK Biobank, including genetic markers, covariates and MRI images are available through application (http://www.ukbiobank.ac.uk/register-apply/). This research has been conducted using the UKB resource under application no. 88320. We are grateful for all the participants in that resource. This manuscript reflects the views of the authors and may not reflect the opinions or views of the UK Biobank funders and investigators.

## Code availability

Software for the PGS pipeline is available at https://github.com/SaraBecelaere/STREAM-PRS. Code for training the genetic algorithm is available at https://github.com/mm-yuan/optimize_phenotyping.

## Author contributions

M.Y. Formal analysis, Methodology, Investigation, Visualization, Data curation, Software, Writing

– original draft, Writing – review & editing

S.G. Investigation, Data curation, Writing – review & editing

N.C. Investigation, Data curation, Writing – review & editing

J.D. Investigation, Data curation, Writing – review & editing

S.B. Investigation, Software, Resources, Writing – review & editing

I.C. Investigation, Software, Resources, Writing – review & editing

P.C. Conceptualization, Methodology, Investigation, Supervision, Funding acquisition, Writing – review & editing, Project administration

## Supporting information

**Supplementary File 1 -Table.xlsx**

Supplementary Table S1: Source data for Fig. 2

Supplementary Table S2: Source data for Fig. 3

**Supplementary File 2 – Additional Materials**

The facial template, nasal landmark labels, and visualizations of all nasal traits are available online at https://doi.org/10.6084/m9.figshare.29621687.

**Supplementary File 3 – Implementation details**

